# Integration of vestibular and hindlimb inputs by vestibular nucleus neurons: Multisensory influences on postural control

**DOI:** 10.1101/661736

**Authors:** Derek M. Miller, Carey D. Balaban, Andrew A. McCall

## Abstract

1.

We recently demonstrated in both decerebrate and conscious cat preparations that hindlimb somatosensory inputs converge with vestibular afferent input onto neurons in multiple CNS locations that participate in balance control. While it is known that head position and limb state modulate postural reflexes, presumably through both vestibulospinal and reticulospinal pathways, the combined influence of the two inputs on the activity of neurons in these brainstem regions is unknown. In the present study, we evaluated the responses of vestibular nucleus (VN) neurons to vestibular and hindlimb stimuli delivered separately and together in conscious cats. We hypothesized that VN neuronal firing during activation of vestibular and limb proprioceptive inputs would be well-fit by an additive model. Extracellular single-unit recordings were obtained from neurons in the caudal aspects of the VN. Sinusoidal whole-body rotation in the roll plane was used as the search stimulus. Units responding to the search stimulus were tested for their responses to 10° ramp-and-hold roll body rotation, 10° extension hindlimb movement, and both movements delivered simultaneously. Composite response histograms were fit by a model of low and high pass filtered limb and body position signals using least squares nonlinear regression. We found that VN neuronal activity during combined vestibular and hindlimb proprioceptive stimulation in the conscious cat is well-fit by a simple additive model for signals with similar temporal dynamics. The mean R^2^ value for goodness of fit across all units was 0.74 ± 0.17. It is likely that VN neurons that exhibit these integrative properties participate in adjusting vestibulospinal outflow in response to limb state.

**New and Noteworthy:** Vestibular nucleus neurons receive convergent information from hindlimb somatosensory inputs and vestibular inputs. In this study, extracellular single unit recordings of vestibular nucleus neurons during conditions of passively applied limb movement, passive whole-body rotations, and combined stimulation, were well fit by an additive model. The integration of hindlimb somatosensory inputs with vestibular inputs at the first stage of vestibular processing suggests vestibular nucleus neurons account for limb position in determining vestibulospinal responses to postural perturbations.

## 1. Introduction

The maintenance of balance is an inherently multimodal process. The central nervous system integrates information from vestibular [1–4], visual [5–8], and proprioceptive [4, 9–12] sensors to generate an estimate of body position and movement in space and to shape corrective postural responses to perturbations [13, 14]. While there are multiple sites in the brainstem that process inputs from these sensory systems, the vestibular nuclei (VN) are of particular importance because they function as key sensory integrators that govern many reflexes relevant to the maintenance of balance, such as vestibulo-ocular and vestibulospinal reflexes [15]. While it has long been appreciated that VN neurons are the site of first synapse of most peripheral vestibular afferents and receive strong proprioceptive projections from the neck, recent work has demonstrated that proprioceptive inputs from the limbs influence VN neuronal firing as well [16, 17].

In situations when neurons receive convergent input from multiple sensory modalities, they must perform a computation and generate a response. This process is called “multisensory integration.” Multisensory integration sometimes results in an output that represents a straightforward summation of responses [18]. In other circumstances, multisensory integration (and in some situations of repeated sensory stimulation or during voluntary movement) responses can be dramatically attenuated or “gated.” [18–21]. In yet other circumstances, responses may be enhanced well beyond those seen when one input is activated alone [22–24]. Previous work in the decerebrate cat model has shown that many vestibular nucleus neurons receiving convergent vestibular afferent and neck proprioceptive afferent inputs integrate those inputs in an additive fashion. Response properties of the individual components (opposing vector orientations, similar sensitivity and phase dynamics) are well-suited to sum and therefore cancel out when coactivated, which has been shown in decerebrate cats during head-on-body movements which simultaneously activate both receptors [25]. Similar findings have been confirmed in some species of conscious non-human primates [26], but not others [27].

Our objective was to determine how VN neurons integrate convergent vestibular and limb proprioceptive inputs. The processing of inputs from these sources is perhaps more complex than those from vestibular and neck proprioceptive inputs. Coactivation of vestibular and neck proprioceptive inputs would be expected to be tightly coupled because, by necessity, nearly all head movements during ordinary activity will result in coactivation of vestibular and neck proprioceptive inputs. A similar tight coupling of vestibular and limb proprioceptive inputs would be expected in some circumstances, such as during ambulation, because head movements are patterned with the step cycle [28]. In contrast, other movements of the weight-bearing limbs would not be expected to have a predictable or appreciable effect on head movement, such as standing in place and raising one leg. Therefore, the processing of proprioceptive information from different areas of the body may be handled differently in the vestibular nuclei. We hypothesized that VN neuronal firing during activation of vestibular and limb proprioceptive inputs would be well-fit by an additive model.

Here, we report results of in vivo recordings from single VN neurons in conscious cats in response to whole-body trapezoid rotation in the roll plane (vestibular stimulation), hindlimb movement in the sagittal plane (limb proprioceptive stimulation), and during simultaneous delivery of both vestibular and limb stimuli. For each neuron, composite response histograms were generated and fit by a model of low and high pass filtered limb and vestibular signals using least squares nonlinear regression. We found that VN neuronal firing was explained by an additive model that incorporated subcomponents of the stimuli. We speculate that VN neurons that exhibit these integrative properties participate in adjusting vestibulospinal outflow in response to limb state.

## 2. Methods

Experimental protocols were reviewed and approved by the University of Pittsburgh’s Institutional Animal Care and Use Committee and conformed to the National Research Council Guide for the Care and Use of Laboratory Animals (National Academies Press, Washington, DC, 2011). Data were collected from four purpose-bred spayed female cats obtained from Liberty Research (Waverly, NY, USA) that were instrumented for chronic single unit recordings using procedures described in previous publications [29–32]. These procedures will be summarized succinctly in the text below.

### 2.1. Surgical Procedures

After a period of acclimation for restraint, each animal underwent a recovery surgery conducted under aseptic conditions in a dedicated operating suite. Animals were initially anesthetized using an intramuscular injection of ketamine (20 mg/kg) and acepromazine (0.2 mg/kg). Animals were intubated, and anesthesia was maintained using isoflurane (1-2%) vaporized in oxygen. An intravenous catheter was inserted in the forelimb and saline was infused intravenously to address fluid loss. A heating pad and an infrared heat lamp were used to maintain the core temperature between 36-38 °C. A midline 1cm craniotomy was made in the posterior aspect of the skull centered over the vestibular nuclei using stereotactic coordinates. A recording chamber (David Kopf Instruments, Tujunga, CA, USA) was centered over the craniotomy and anchored to the skull using stainless steel screws and Palacos bone cement (Zimmer, Warsaw, IN, USA). Rostral to the recording chamber, a fixation plate was mounted to the skull in a similar fashion. Electromyographic (EMG) patch electrodes were sutured to the biceps femoris and vastus lateralis muscles of both limbs through incisions on the lateral aspect of the thigh; leads were routed subcutaneously to a connector mounted on the skull. One of the four animals had significant bleeding from the cerebellum at the time of craniotomy requiring repeated application of hemostatic agents; after the surgery, the animal was posturally unsteady for several weeks. Postmortem histological examination demonstrated partial degeneration of the cerebellar cortex ipsilateral to the recording site. Despite this injury, the single unit data from the VN of this animal were indistinguishable from those obtained from the other three animals and thus were included in this report. Post-surgical analgesia was provided continuously for 72 hours through a transdermal fentanyl delivery system (25 μg/h, Janssen Pharmaceutical Products, Titusville, NJ, USA). Amoxicillin (50 mg, BID) was administered orally for ten days following surgery.

### 2.2. Vestibular Stimulation and Hindlimb Movement Protocols

Animals were acclimated over a period of several weeks to head and body restraint while table movements and hindlimb ramp-and-hold movements were applied. The animal was placed in a modified restraint bag (Four Flags Over Aspen, Aspen, CO, USA) with a hole cut in the rear to allow access to the hindlimb. The hindlimb was secured via a Velcro strap placed at the ankle to a servo-controlled motor capable of delivering movements in the rostral-caudal axis (i.e. hindlimb extension and flexion). The torso was fit snugly into a padded cylindrical tube secured to the stereotaxic frame, and straps were placed around the animal’s body to snugly secure the animal. The head was immobilized by attaching the fixation plate to a post mounted on the stereotaxic frame. The stereotaxic frame was mounted atop a servo-controlled hydraulic tilt table (NeuroKinetics, Pittsburgh, PA, USA) capable of delivering movements in the pitch and roll planes (Fig. 1).

**Figure 1.**
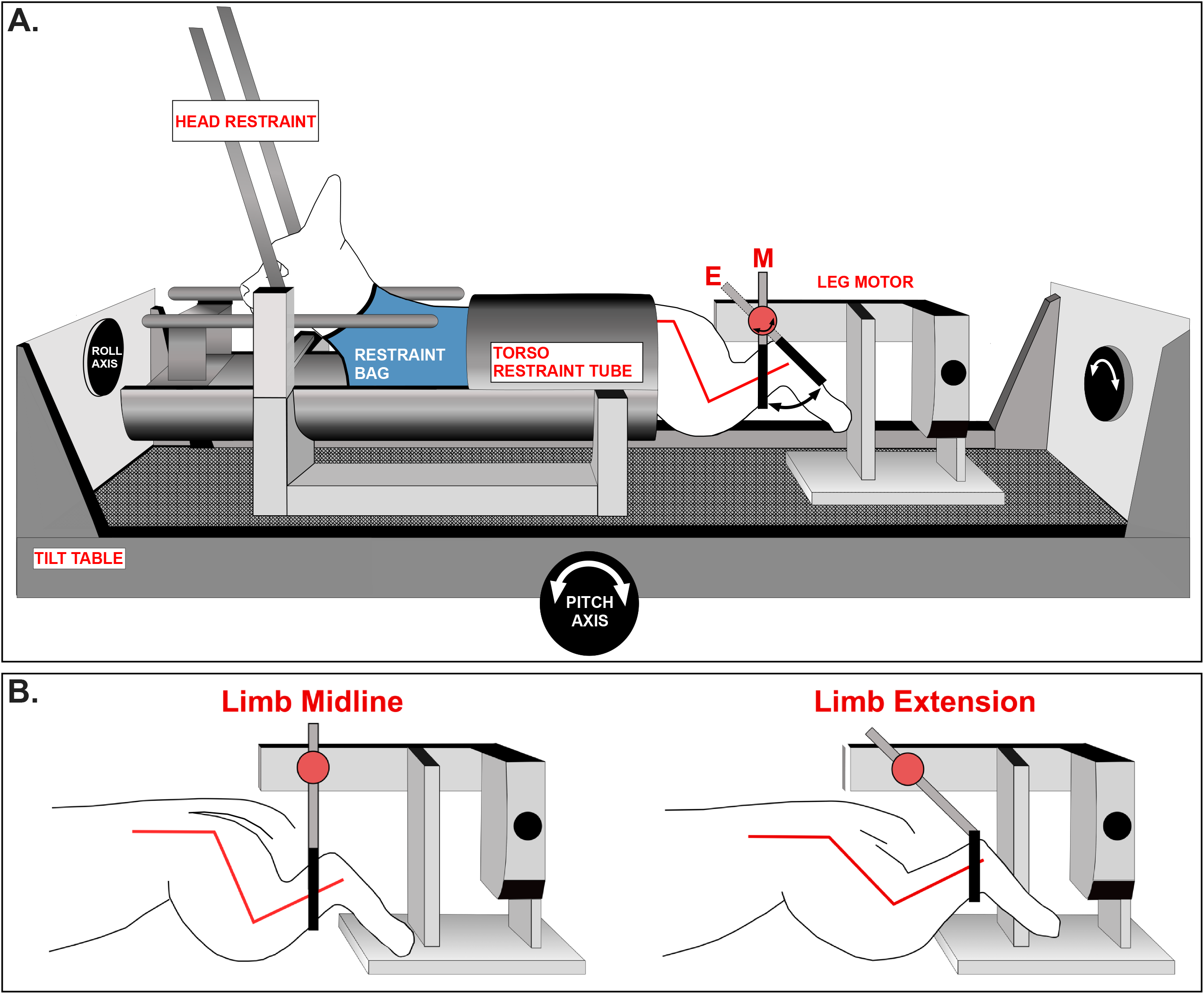
**A.** Awake felines were placed into a restraint device mounted on a servo-controlled tilt table. The two-axis tilt table was used to provide vestibular stimulation by rotating the animal in the roll plane. **B.** A servo-controlled motor positioned behind the animal provided hindlimb movement. The hindlimb was placed through a hole in the restraint bag and secured to the servo-controlled motor via a Velcro strap attached immediately proximal to the ankle. Hindlimb position at midline and extension, with relative angles of the knee and hip joints, are shown. Figure modified and reproduced with permission from Springer, *Miller et al., 2017*.

Sinusoidal whole-body rotation in the roll plane was used as the search stimulus to identify neurons receiving vestibular afferent input (Fig. 2). Units responding to the search stimulus were evaluated further for responses during three conditions in succession (hereafter referred to as integration trials): (1) ramp-and-hold body roll rotation; (2) ramp-and-hold leg extension; and (3) simultaneously delivered ramp-and- hold body roll and leg extension (Fig. 3). Maximum displacement during the hold segment of body roll and leg extension was 10° and 60°, respectively. Each trial incorporating all three conditions was 18 seconds in duration with each ramp phase lasting one second and each hold phase lasting two seconds. In most instances, ten trials were repeated for each isolated neuron. In some instances, a given unit was lost prior to completion of all trials; neurons with fewer than five trials were excluded from analysis. During most trials, animals permitted passive hindlimb movement, as there was no change in hindlimb muscle EMG activity, and the limb movement recorded from a potentiometer on the servomotor was identical to the command signal provided by the computer controlling the servomotor. Trials in which EMG activity increased over baseline during the hindlimb movement or where the hindlimb movement deviated from that delivered by the servomotor were excluded from the analyses described below.

**Figure 2.**
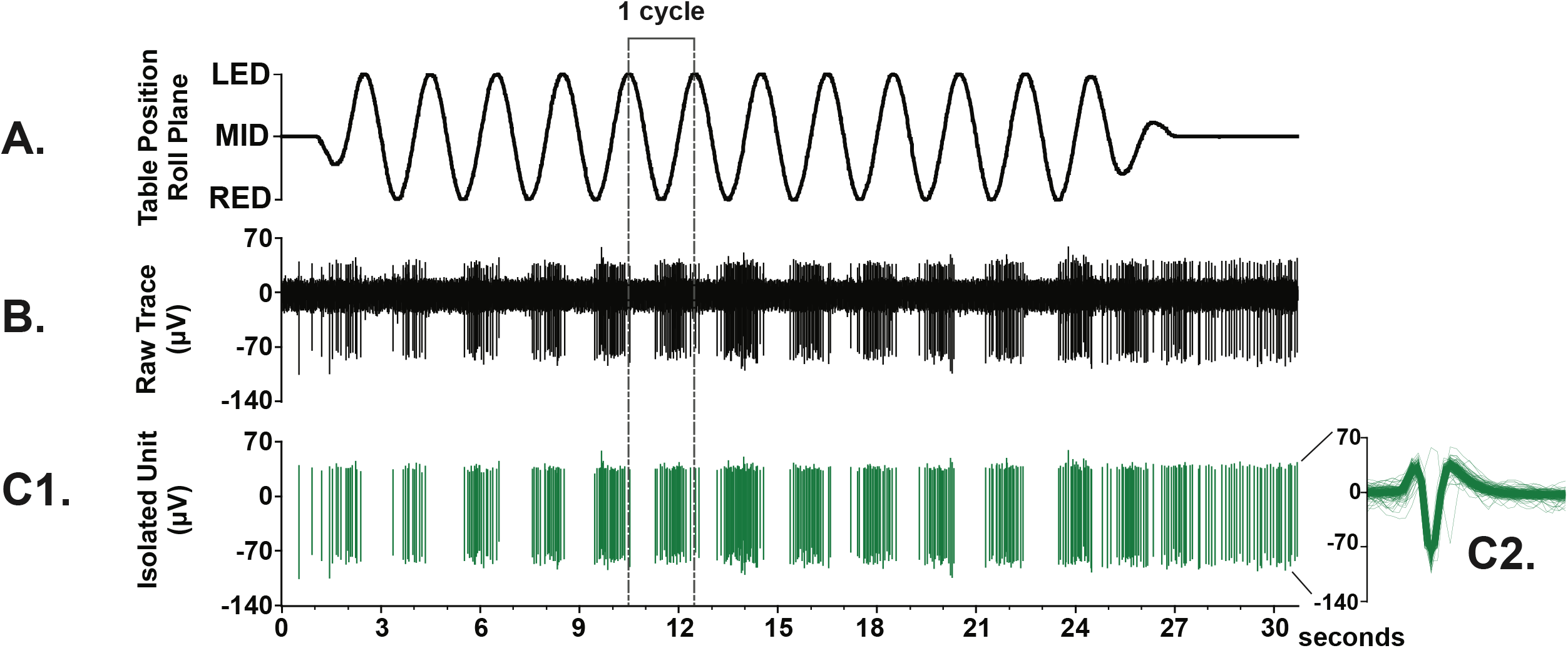
Example of a single vestibular nucleus (VN) neuron whose activity was robustly modulated by the search stimulus. **A.** Sinusoidal whole-body rotation in the roll plane was used as the search stimulus to identify neurons that received vestibular inputs (*tilt frequency, 0.5Hz; tilt amplitude, 5°*). **B.** Raw neural data sampled at 25 kHz. **C1.** Tracing showing the isolated neuron of interest. **C2.** The overlaid spike-sorted waveforms were equivalent across all the trial, indicating that activity was sampled from the same neuron throughout the recording.

**Figure 3.**
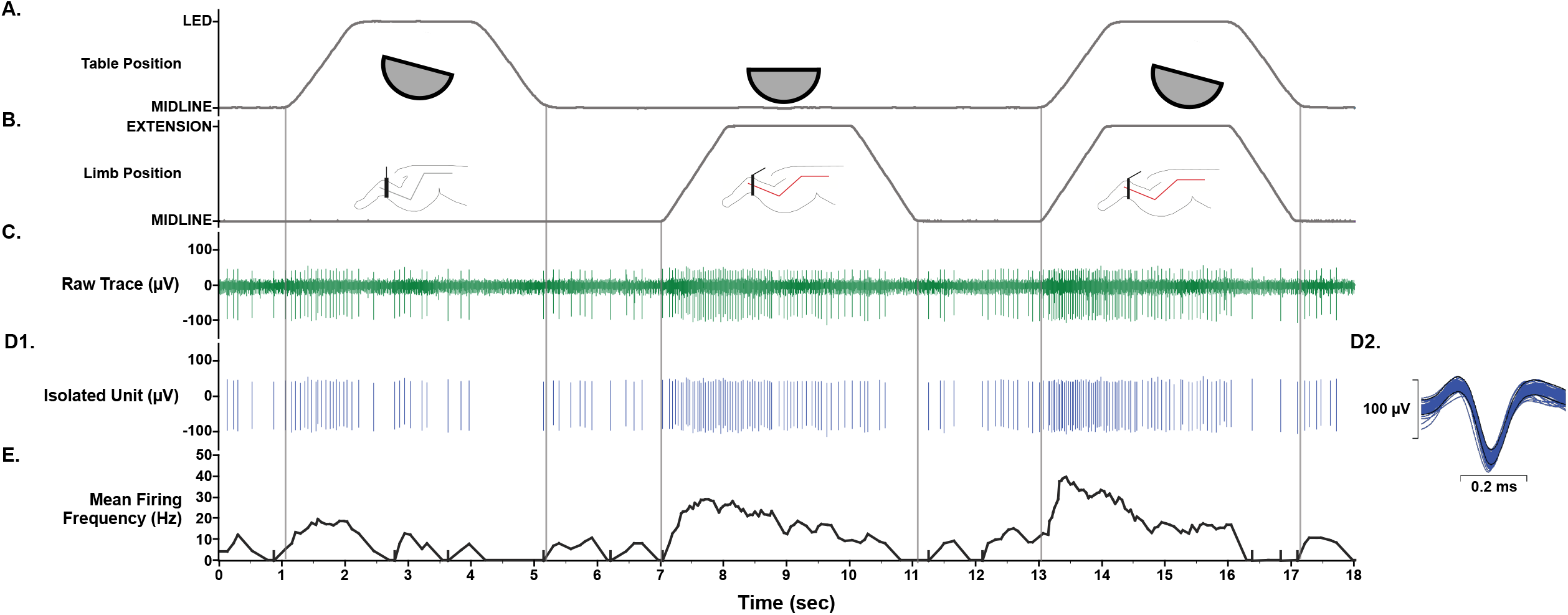
Units responding to the vestibular search stimulus were further tested for their responsiveness to (**A**) 10° ramp-and-hold roll body and head rotation, (**B**) 60° ramp-and-hold hindlimb extension movement, and both movements delivered simultaneously. **C.** Raw neural data was sampled at 25 kHz. **D1**. Unit activity was isolated using template-matching and response histograms were generated and fit by a model of low and high pass filtered limb & body position and velocity signals using least squares nonlinear regression. **D2**. The average spike-sorted waveforms (insets on right) were equivalent across the trial, indicating that activity was sampled from the same neuron throughout the recordings. E. Mean firing frequency (*bin width, 0.25 sec*).

### 2.3. Neural Recording Procedures

Extracellular recordings were made from VN neurons using 4-6 MΩ epoxy-insulated tungsten micro-electrodes (Frederick Haer, Bowdoin, ME, USA). An XY positioner (David Kopf Model 608B) was secured atop the recording chamber, and the electrode was introduced via a 25-gauge stainless steel guide tube inserted through the dura and into the cerebellum. The electrode was lowered into the medulla using a hydraulic microdrive (David Kopf Model 650). Neural activity was amplified by a factor of 1K or 10K, bandpass filtered 0.3 - 10 kHz, and sampled at 25 kHz using a Cambridge Electronic Design Micro 1401 mk2 data collection system and Spike 2 version 8 software (Cambridge, UK). EMG data from the hindlimb musculature were amplified by 1000, bandpass filtered 0.01 - 10 kHz and sampled at 1 kHz. Signals from potentiometers on the tilt table and servo-controlled leg motor provided body and hindlimb position, respectively, and were each sampled at 100 Hz.

### 2.4. Vestibular and Limb Signal Identification

Spike occurrences were binned at 10 msec intervals and converted into instantaneous spike rates by dividing by the number of stimulus cycles and multiplying by 100. The table and limb position traces were centered at zero. The table and limb velocities were calculated with the ‘gradient.m’ function in MATLAB (The MathWorks, Natick, MA) at the 100 Hz sample rate. The limb position, limb velocity, table roll position, table roll velocity and unit response data were then decimated to a 20 Hz rate using the MATLAB ‘decimate.m’ function for model estimation. The first step in analysis was generation of a formal, parametric description of dynamic signal processing by these units. The MATLAB function ‘lsim.m’ was used to implement low pass and high pass representations of the limb and table stimulus position and velocity traces, with simple first order transfer functions of 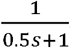 and 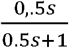, respectively. The spike rate data sets were then fitted as a weighted linear sum of a baseline firing rate and direction sensitivities for the low pass (LP) and high pass (HP) roll position, HP roll velocity, LP and HP limb position and LP and HP pass limb velocity, with sensitivities estimated by the MATLAB ‘lsqnonlin.m’ function (Levenberg – Marquardt nonlinear least squares optimization). These models assumed additivity of all signals for the entire stimulus presentation period (roll alone, limb alone, then roll plus limb). Further statistical analysis was conducted in either SPSS version 25 (*with R and Python extensions*) or in MATLAB.

#### 2.4.1. First Iteration

An iterative strategy was followed to identify an uncorrelated set of vestibular and proprioceptive signals as linear combinations of the initial set of 13 signal components. The initial optimization step produced parameter estimates for:

1. baseline activity (a constant)
2. HP limb position sensitivity for extension
3. HP limb position sensitivity for return to midline (hereafter “flexion”)
4. HP limb velocity sensitivity for extension
5. HP limb velocity sensitivity for flexion
6. LP roll position sensitivity for ipsilateral ear up rotation
7. LP roll position sensitivity for ipsilateral ear down rotation
8. HP roll position sensitivity for ipsilateral ear up rotation
9. HP roll position sensitivity for ipsilateral ear down rotation
10. LP limb position sensitivity for extension
11. LP limb position sensitivity for flexion
12. HP roll velocity sensitivity for ipsilateral ear up rotation
13. HP roll velocity sensitivity for ipsilateral ear down rotation

Exploratory examination of the scatterplot matrix of these parameters for all units, validated by linear regression analysis, showed the following strong linear dependencies between components:

1. The high pass roll position sensitivities in the ipsilateral ear down and ipsilateral ear up sensitivities were highly correlated (*adjusted R^2^ = 0.959*). The linear relationship was:

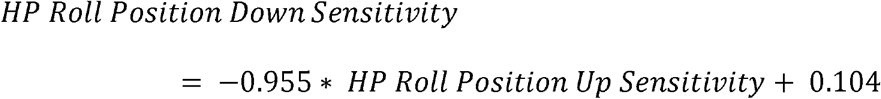 Given the high R^2^, the negligible intercept parameter, and a slope that does not differ from −1, high pass roll position was represented as a fully rectified process with a single sensitivity parameter for the second iteration.
2. The LP roll position sensitivities in the ipsilateral ear down and ipsilateral ear up sensitivities were highly correlated (*adjusted R^2^ = 0.910*). The linear relationship was given by:

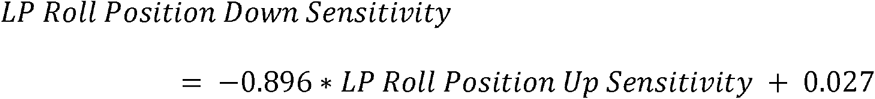 Because the R^2^ is very high, the slope that does not differ from −1 and the estimated intercept is negligible, low pass roll position was represented as a fully rectified process with a single sensitivity parameter for the second iteration.
3. The LP limb position for flexion and extension sensitivities were highly correlated (*adjusted R^2^ = 0.908*). The linear relationship was described by:

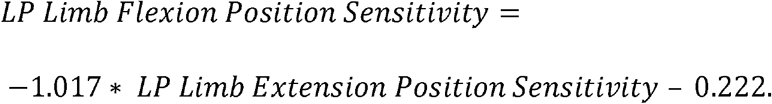 As in the case of low pass roll position, the R^2^ is very high, the slope that does not differ from −1 and the estimated intercept is negligible. Hence, low pass limb position was represented as a fully rectified process with a single sensitivity parameter for the second iteration
4. The high pass roll velocity sensitivities in the ipsilateral ear down and ipsilateral ear up sensitivities were correlated significantly, but to a lesser degree (*adjusted R^2^ = 0.437*). The linear relationship was:

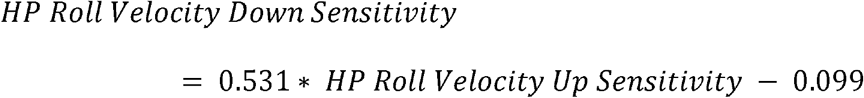

As a result, a single unrectified high pass roll velocity sensitivity process was implemented in the next iteration of the model.

#### 2.4.2. Second Iteration

The **second iteration** began with a new least squares estimation of parameters for representing the unit responses as a weighted sum of nine components (with the same dynamics as described previously):

1. baseline activity (constant)
2. HP limb position sensitivity for extension
3. HP limb position sensitivity for flexion
4. HP limb velocity sensitivity for extension
5. HP limb velocity sensitivity for flexion
6. rectified HP roll position sensitivity
7. HP roll velocity sensitivity (unrectified, ipsilateral ear-up positive)
8. rectified LP limb position sensitivity
9. rectified LP roll position sensitivity

Graphical (visualization) and linear regression analysis revealed the sensitivities of the high pass roll velocity (unrectified) and rectified high pass roll position processes are highly correlated (*adjusted R^2^=0.674*), with the sensitivity of the former equal to 0.291 times the latter with no constant term.

#### 2.4.3. Third Iteration

For the **third iteration**, the model components were:

1. baseline activity (constant)
2. HP limb position sensitivity for extension
3. HP limb position sensitivity for flexion
4. HP limb velocity sensitivity for extension
5. HP limb velocity sensitivity for flexion
6. rectified HP roll position + 0.291* HP roll velocity sensitivity (ipsilateral ear-up positive)
7. residual HP roll velocity sensitivity (unrectified, ipsilateral positive)
8. rectified LP limb position sensitivity
9. rectified LP roll position sensitivity Analysis of relationships among the fitted parameters for this modified model showed that the baseline activity was related linearly (*adjusted R^2^=0.604*) to the rectified low pass limb position sensitivity as 23.441-5.163 * rectified low pass limb position sensitivity. This was incorporated in a **fourth iteration**.

#### 2.4.4. Fourth Iteration

For the **fourth iteration**, the model components were:

1. new baseline parameter (constant-5.163 *rectified LP limb position sensitivity)
2. HP limb position sensitivity for extension
3. HP limb position sensitivity for flexion
4. HP limb velocity sensitivity for extension
5. HP limb velocity sensitivity for flexion
6. rectified HP roll position sensitivity + 0.291* HP roll velocity sensitivity (ipsilateral ear-up positive)
7. residual HP roll velocity sensitivity (unrectified, ipsilateral positive)
8. rectified LP limb position sensitivity −5.68 sp/s (a constant to adjust its set point)
9. rectified LP roll position sensitivity

The analysis of the parameter estimates from this final iteration showed one significant relationship (*adjusted R^2^=0.36*) for the high pass limb velocity extension and flexion sensitivities. The latter sensitivity was equal to 0.402 (±0.075) times the former.

#### 2.4.5. Fifth Iteration

The **final form of the model for functional unit analysis** was the weighted sum of nine component signals:

1. baseline activity (constant-5.1635.163 * rectified LP limb position sensitivity),
2. HP limb position sensitivity for extension
3. HP limb position sensitivity for flexion
4. HP limb velocity sensitivity for extension (residual)
5. HP limb velocity sensitivity for flexion– 0.402* HP limb extension velocity sensitivity
6. rectified HP roll position sensitivity + 0.291* HP roll velocity sensitivity (ipsilateral ear-up positive)
7. residual HP roll velocity sensitivity (unrectified, ipsilateral positive)
8. rectified LP limb position sensitivity −5.68 sp/s (a constant to adjust its set point)
9. rectified LP roll position sensitivity

### 2.5. Histologic Analysis

After the completion of data collection from each animal, two electrolytic lesions were made near the recording sites in the brainstem by passing a 0.5-1 mA current through a low impedance (0.5 MΩ) electrode for up to 60 seconds. Electrolytic lesions were allowed to mature for seven days prior to euthanasia. Animals were deeply sedated with an initial intramuscular injection of ketamine (20 mg/kg) and acepromazine (0.2 mg/kg) followed by an intraperitoneal injection of pentobarbital sodium (40 mg/kg) and then perfused transcardially with 10% formalin. The brainstem was removed, fixed in 10% formalin, embedded in 2% agar, and cut transversely at a thickness of 50 μm using a freezing microtome. Sections were stained with 1% thionine. Recording locations were reconstructed with reference to the electrolytic lesions, the relative positions of the recording tracks, and the relative depths of the units.

## 3. Results

Complete data were obtained from 129 VN units of four conscious felines. Composite response histograms were generated for each unit, and the response histograms were fit by an additive model of low and high pass filtered limb and body position signals using least squares non-linear regression as outlined above (Fig. 4). This additive model for roll and limb stimulation fit the population of units with high fidelity. The mean R^2^ value for goodness of fit across all units was 0.74 ± 0.17 (range, 0.31 - 0.98). The distribution of the R^2^ values is shown in figure 5 and was negatively skewed (skew statistic, −0.484).

**Figure 4.**
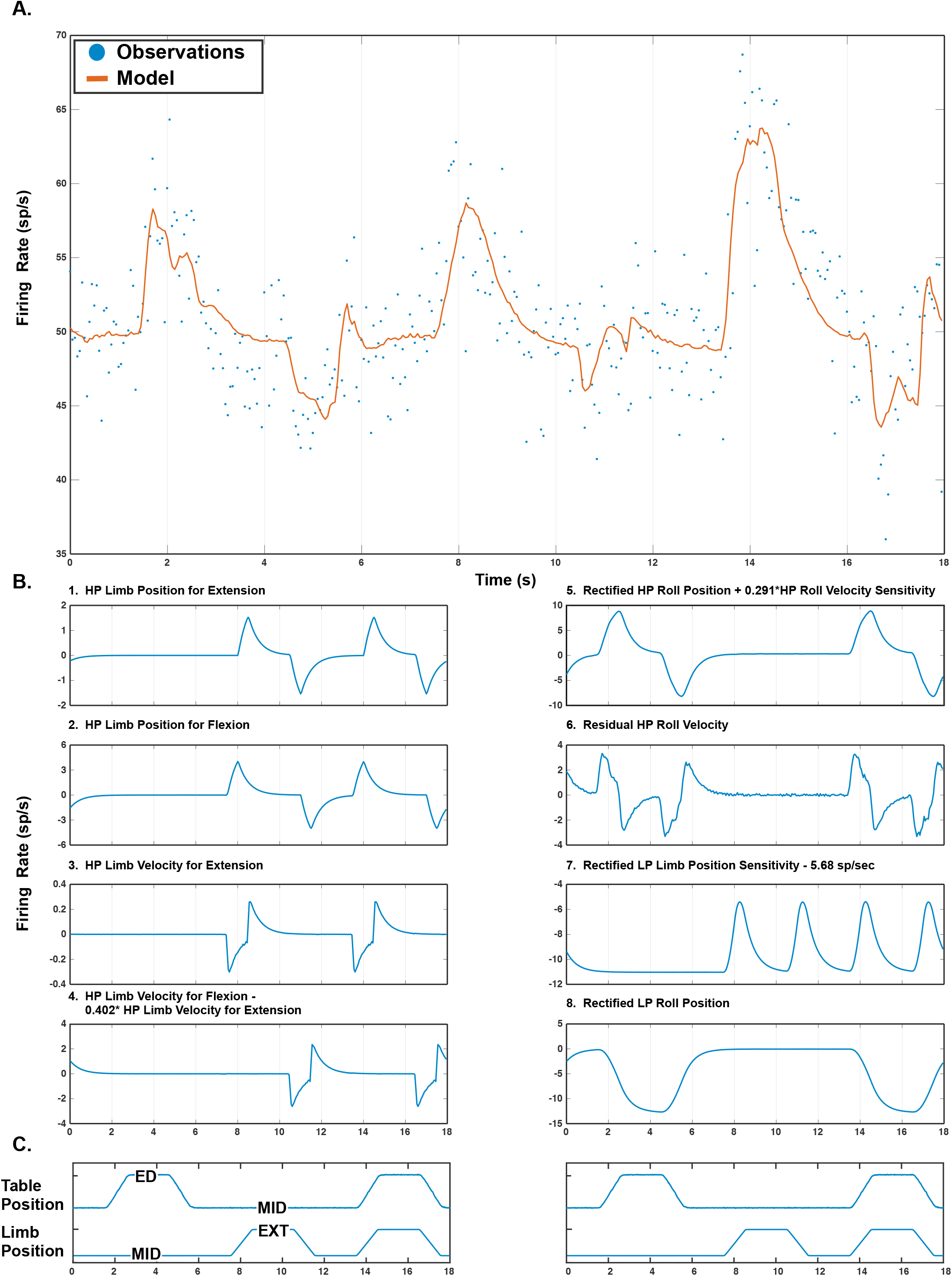
An example response of a VN neuron to roll body rotation, hindlimb movement and both movements delivered simultaneously. The response to simultaneous delivery of roll and limb stimuli results from the weighted sum of the individual components. **A**. The observed firing rate (blue dots) and the modeled firing rate (orange solid line, see section 3.4) are shown. **B**. The individual signal components for the unit in (**A**) are plotted (with the exception of baseline firing rate, which is a flat line). **C**. Limb and table position. Abbreviations: *ED, ear down; EXT, extension; MID, midline*.

**Figure 5.**
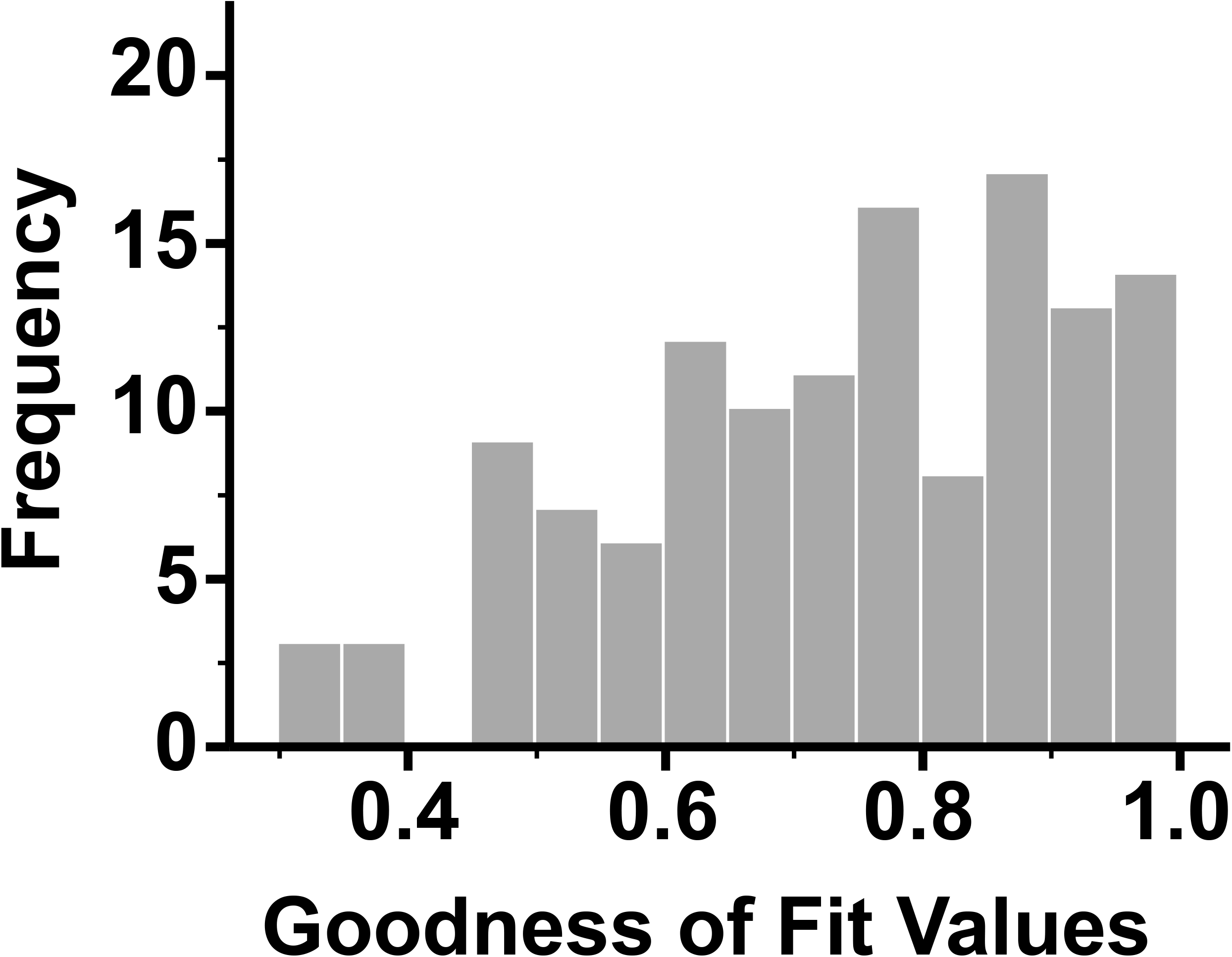
A histogram of the distribution of R^2^ values for the fit of an additive model of low and high pass filtered limb and table movement dynamics (indicative of limb proprioceptive and vestibular signals, respectively) using least squares nonlinear regression to neural firing of individual VN neurons.

### 3.1. Vestibular Dynamics

The vestibular dynamics are represented robustly by signal comprised of a <u>linear combination of the rectified high pass roll position and velocity subcomponents; the sensitivity to this variable will hereafter be referred to as “vestibular sensitivity</u>.” The signal was identified through a step-wise elimination of the partial correlations of covarying components, and thus carries a substantial amount of the vestibular sensitivity. All cells in our dataset had non-zero estimates for this parameter. Note that parameters representing other vestibular dynamics were also in the final model; those parameters followed similar distributions as shown for this variable.

We investigated the magnitude of the vestibular responses, irrespective of polarity or directionality, by taking the absolute value of the sensitivity of this component. The distribution of the absolute sensitivity is shown in figure 6. Evaluation of the histogram in figure 6b reveals an exponential distribution with a scale parameter (σ) of 0.390. We divided the distribution into three distinct sensitivity classes based on the reciprocal of the scale parameter (Table 1):

> **Low** absolute vestibular sensitivity ≤ 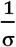; 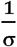 < **intermediate** vestibular sensitivity ≤ 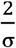; **high** absolute vestibular sensitivity > 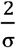

**Figure 6.**
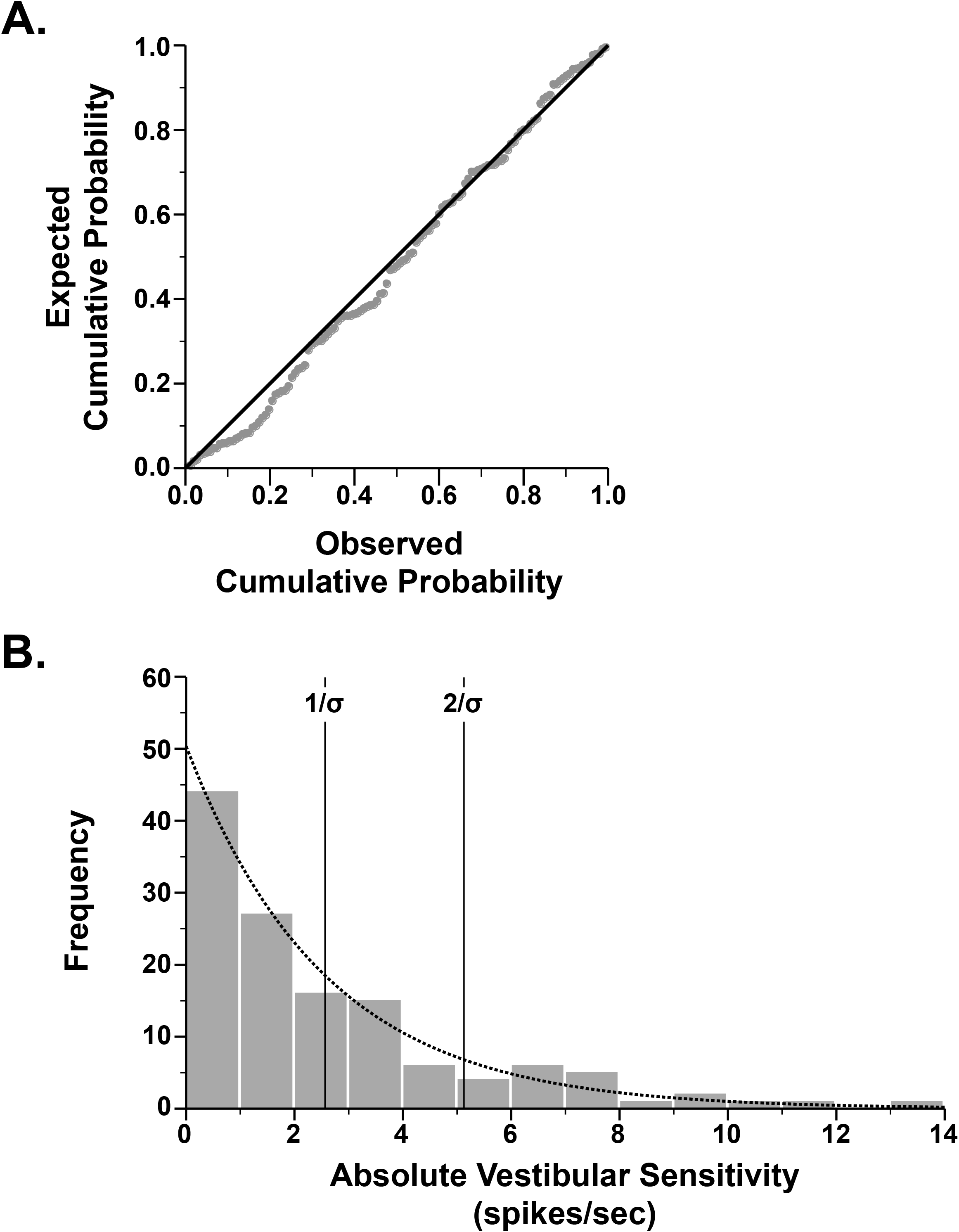
**A.** Exponential probability-probability (P-P) plot of two cumulative distribution functions. The expected cumulative probability for an exponential distribution (y) is plotted against the observed cumulative probability (x) for the absolute value of the vestibular sensitivity variable. **B.** A histogram of the distribution of the absolute value of the vestibular sensitivity variable is positively skewed. It is fit-best by an exponential distribution with a scale parameter (σ) of 0.390. The units were divided into distinct classes on the basis of the reciprocal of the scale parameter and were classified as having either low absolute vestibular sensitivity, intermediate absolute vestibular sensitivity, or high absolute vestibular sensitivity.

**Table 1.**

| **Table 1** | **Subclass** | **Count** | **Mean** | **SD** | **Min** | **Max** |
| --- | --- | --- | --- | --- | --- | --- |
| **Absolute Vestibular Sensitivity** (spikes/sec) | LOW | 82 | 0.969 | 0.741 | 0.006 | 2.538 |
|  | INTERMEDIATE | 27 | 3.507 | 0.628 | 2.625 | 5.067 |
|  | HIGH | 20 | 7.814 | 2.236 | 5.284 | 13.755 |
| **Absolute LP Limb Position Sensitivity** (spikes/sec) | LOW | 86 | 1.256 | 0.853 | 0.002 | 3.170 |
|  | INTERMEDIATE | 27 | 4.541 | 0.993 | 3.279 | 6.340 |
|  | HIGH | 16 | 11.259 | 6.315 | 6.571 | 25.954 |
| **Absolute HP Limb Velocity Sensitivity** (spikes/sec) | LOW | 90 | 0.092 | 0.064 | 0.002 | 0.227 |
|  | INTERMEDIATE | 20 | 0.327 | 0.067 | 0.245 | 0.438 |
|  | HIGH | 19 | 0.764 | 0.238 | 0.475 | 1.371 |

The majority of neurons (63.6%) were classified into the low absolute vestibular sensitivity category (82/129 neurons). The mean sensitivity for these neurons was 0.969 ± 0.741 spikes/sec, and sensitivities ranged from a minimum of 0.006 to a maximum of 2.538 spikes/sec. Approximately twenty-one percent of neurons were classified as having intermediate absolute vestibular sensitivity (20.9%; 27/129 neurons). These neurons had vestibular sensitivities ranging from 2.625 to 5.067 spikes/sec (mean ± SD, 3.507 ± 0.628 spikes/sec). The remaining 15.5% of neurons were classified as having high absolute vestibular sensitivity (20/129 neurons). Vestibular sensitivity values for neurons in this category ranged from 5.284 to 13.755 spikes/sec, and the mean was 7.814 ± 2.236 spikes/sec.

### 3.2. LP Limb Position Dynamics

Dynamics of the response to low pass limb movement are represented well by a <u>LP limb position sensitivity signal</u> (*rectified low pass limb position −5.68* spikes/sec). <u>The sensitivity to this variable will hereafter be referred to as “LP limb position sensitivity</u>.” The signal was identified through a step-wise elimination of the partial correlations of covarying components, and thus carries a substantial amount of the LP limb position sensitivity. All cells in our dataset had non-zero estimates for this parameter. The sensitivities of neuronal responses to this signal showed a similar distribution to that of the vestibular sensitivity signal after taking the absolute value of the limb sensitivities. Evaluation of the histogram in figure 7b reveals an exponential distribution with a scale parameter (σ) of 0.314. The distribution was divided into three distinct classes based on the reciprocal of the scale parameter (Table 1):

> **Low** absolute limb sensitivity ≤ 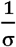; 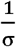 < **intermediate** absolute limb sensitivity ≤ 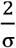; **high** absolute limb sensitivity > 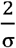

**Figure 7.**
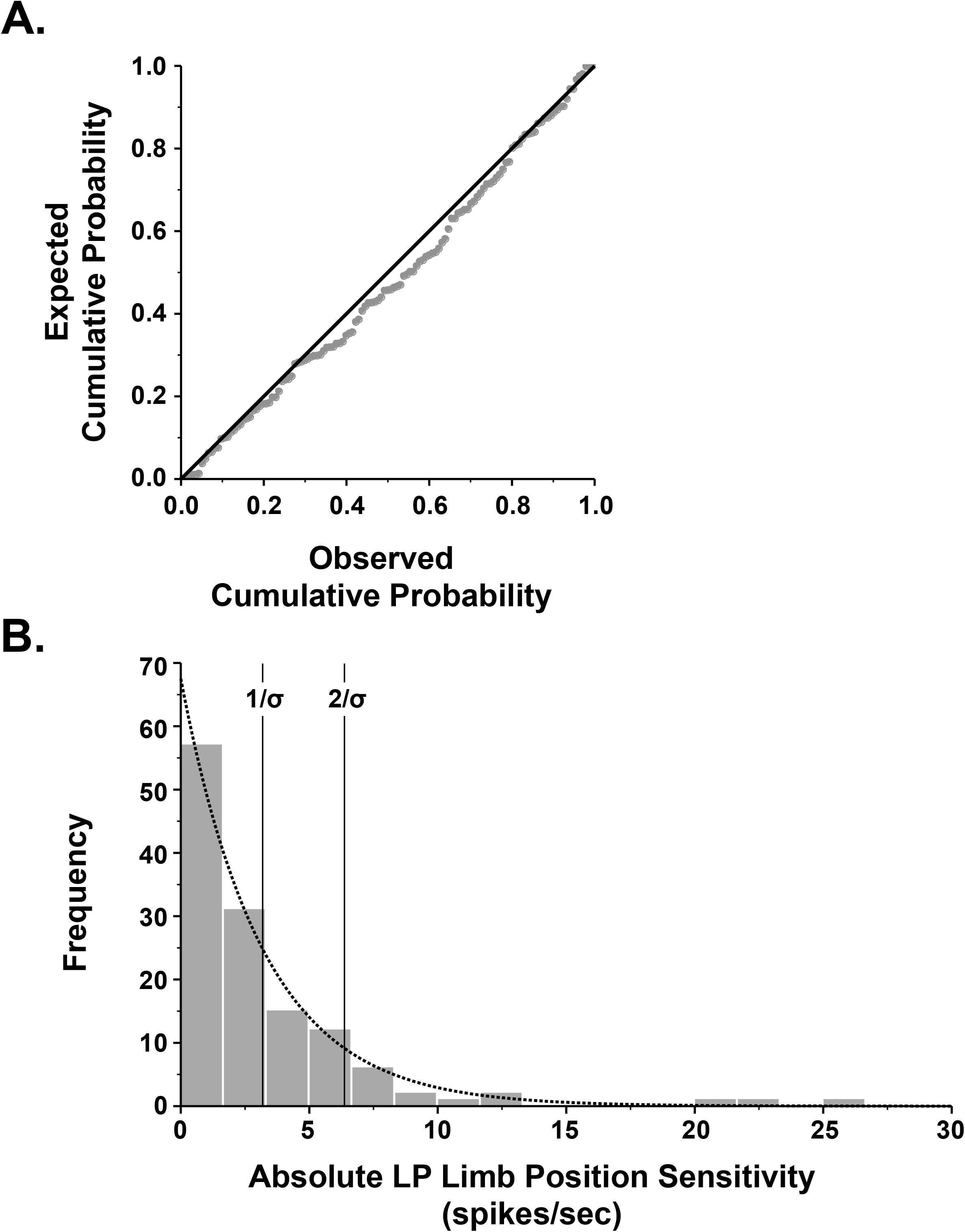
**A.** Exponential P-P plot of the absolute value of LP limb position sensitivity. **B.** A histogram of the distribution of the absolute value of the LP limb position sensitivity is positively skewed and is fit-best by an exponential distribution with a scale parameter (σ) of 0.314. The units were divided into distinct classes on the basis of the reciprocal of the scale parameter and were classified as having either low absolute LP limb position sensitivity, intermediate absolute LP limb position sensitivity, or high absolute LP limb position sensitivity.

Most neurons were classified into the low absolute sensitivity to limb movement category (66.7%, 86/129 neurons). These neurons had sensitivities ranging from 0.002 to 3.170 spikes/sec and had a group mean of 1.256 ± 0.853 spikes/sec. Neurons with intermediate absolute limb sensitivity were the next most common group, representing approximately twenty-one percent (20.9%; 27/129 neurons) of all neurons. The sensitivities of these neurons ranged from a low of 3.279 to a high of 6.340 spikes/sec (mean ± SD, 4.541 ± 0.993 spikes/sec). Finally, neurons in the high sensitivity class were least common (12.4%, 16/129 neurons). The mean sensitivity for this group was approximately 9-fold higher than the mean sensitivity for the low sensitivity neurons and was 2.5-fold higher than intermediate sensitivity neurons (range, 6.571 – 25.954 spikes/sec; mean ± SD, 11.259 ± 6.315 spikes/sec).

### 3.3. Combined Vestibular and LP Limb Position Sensitivities

Neurons could be further divided into nine groups on the basis of vestibular and low pass limb position sensitivities (Fig. 8A; Table 2). The most common unit responses (54/129 units; 41.9%) were neurons with low absolute vestibular and limb sensitivities (Fig. 8B1). Intermediate absolute vestibular – low absolute limb sensitivity neurons (17/129 units; 13.2%; Fig. 8B2) and low absolute vestibular – intermediate absolute limb sensitivity neurons (14/129, 10.9%; Fig. 8B4) were the next most common subcategories. High absolute vestibular – low absolute limb sensitivity neurons were robustly sensitive to vestibular stimulation but were minimally responsive to limb movement; they made up 11.6% (15/129 neurons; Fig. 8B7) of the population. On the opposite end of the spectrum, low absolute vestibular – high absolute limb sensitivity neurons (14/129 neurons; 10.9%; Fig. 8B3) were preferentially responsive to limb movement. The remaining four groups represented 11.6% of the neurons. Intermediate absolute vestibular – intermediate absolute limb sensitivity neurons made up seven percent of the population (9/129 neurons; Fig. 8B5). A single neuron had intermediate absolute vestibular - high absolute limb sensitivity (0.8%; Fig. 8B6). Four neurons (3.1%) were highly sensitivity to vestibular stimulation and had intermediate sensitivity to limb movement (Fig. 8B8). Finally, one neuron was classified as having high absolute vestibular - high absolute limb sensitivity (0.8%; Fig. 8B9).

**Figure 8.**
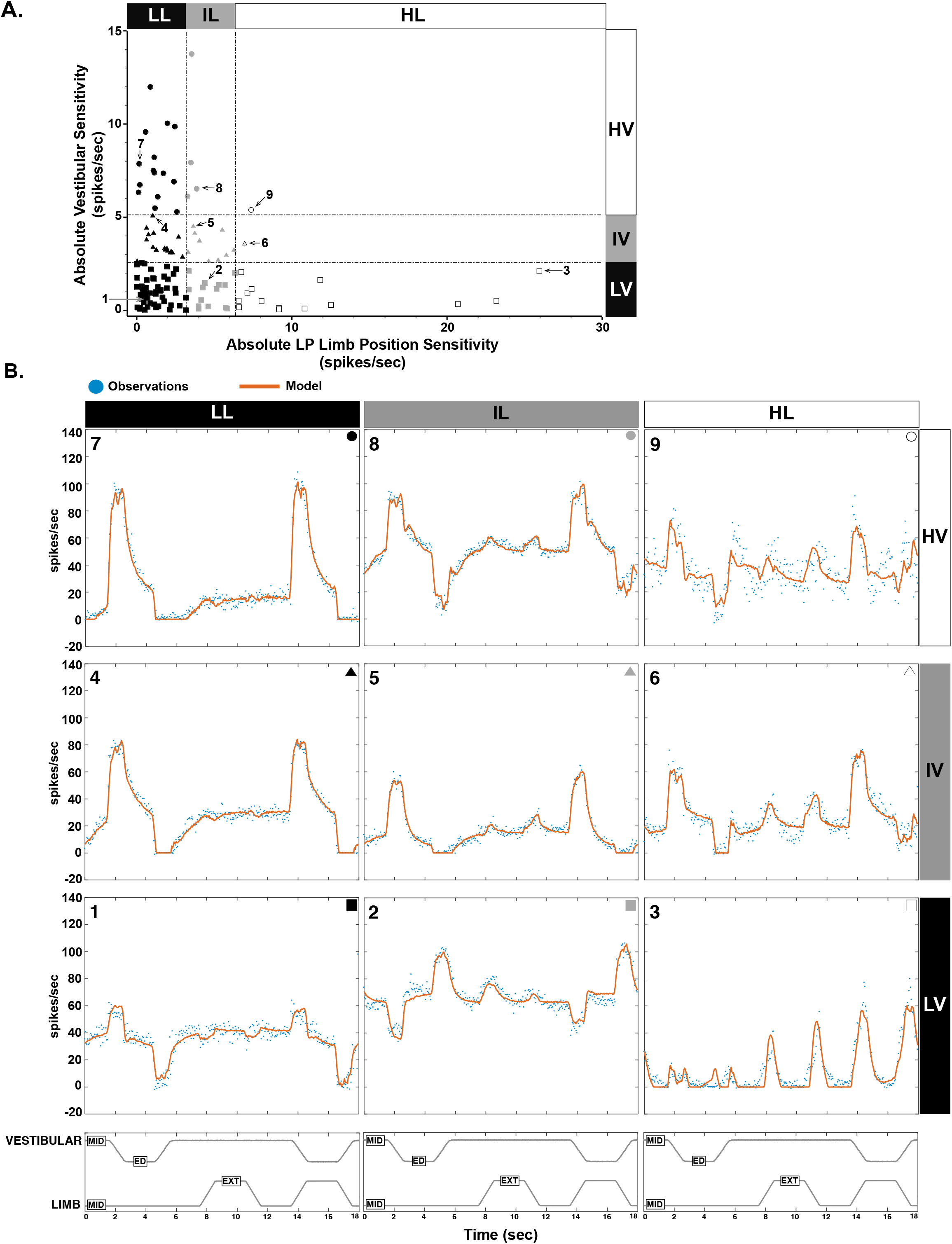
**A.** Neurons could be further divided into nine groups on the basis of vestibular and low pass limb position sensitivities. Units labeled 1 through 9 correspond to the example units shown in B. **B**. The averaged responses of 9 units represent the salient features of VN neuron responses to combined vestibular and hindlimb stimulation. The observed firing rate (blue dots) and the modeled firing rate (the fit of the model to the data; orange solid line, see section 3.4) are shown for each unit. Units 7-9 are classified as having high absolute vestibular sensitivity, units 4-6 have intermediate absolute vestibular sensitivity, and units 1-3 have low absolute vestibular sensitivity. Units 1,4, and 7 have low absolute limb sensitivity, units 2, 5, and 8 have intermediate absolute limb sensitivity, while units 3, 6, and 9 have high absolute limb sensitivity. Abbreviations: *ED, ear down; EXT, extension; MID, midline*

**Table 2.**
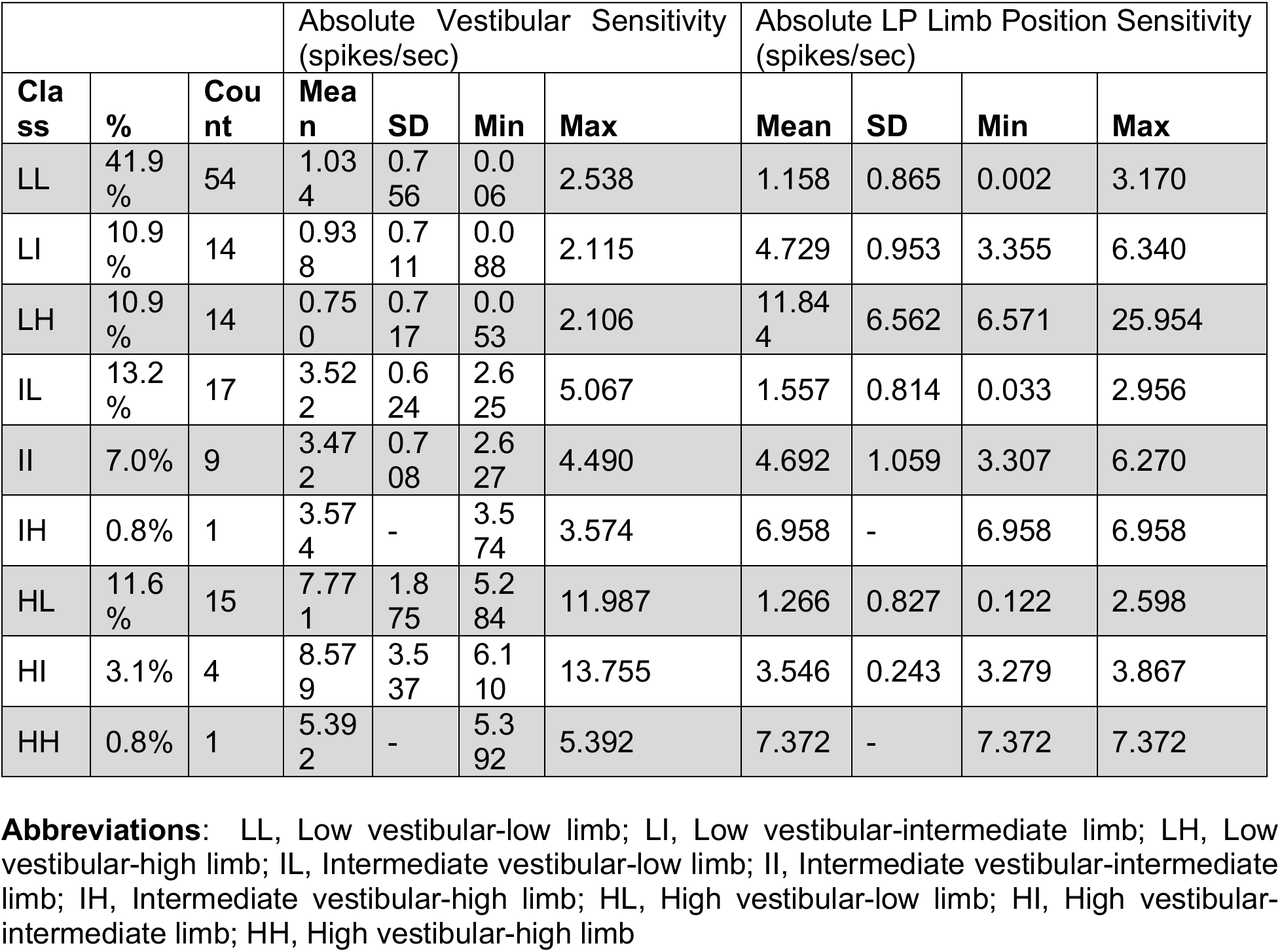

### 3.4. HP Limb Velocity Dynamics

Dynamics of the response to high pass limb movement are represented well by the <u>HP limb velocity sensitivity signal</u> (*HP limb position velocity sensitivity for flexion – 0.402* HP limb extension velocity sensitivity*). <u>The sensitivity to this variable will hereafter be referred to as “HP limb velocity sensitivity</u>.” The signal was identified through a stepwise elimination of the partial correlations of covarying components, and thus carries a substantial amount of the HP limb velocity sensitivity. All cells in our dataset had nonzero estimates for this parameter. Categorization of responses to this signal showed a similar distribution to that of the vestibular and LP limb position sensitivity signals. We investigated the magnitude of the HP limb velocity responses by taking the absolute value of the sensitivity of this component. The distribution of the absolute sensitivity is shown in figure 9. Evaluation of the histogram in figure 9b reveals an exponential distribution with a scale parameter (σ) of 4.401, and the distribution was divided into three distinct sensitivity classes based on the reciprocal of the scale parameter (Table 1):

> **Low** absolute HP limb velocity sensitivity ≤ 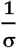; 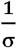 <
>
> **intermediate** HP limb velocity sensitivity ≤ 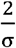; **high** HP limb velocity sensitivity > 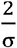

**Figure 9.**
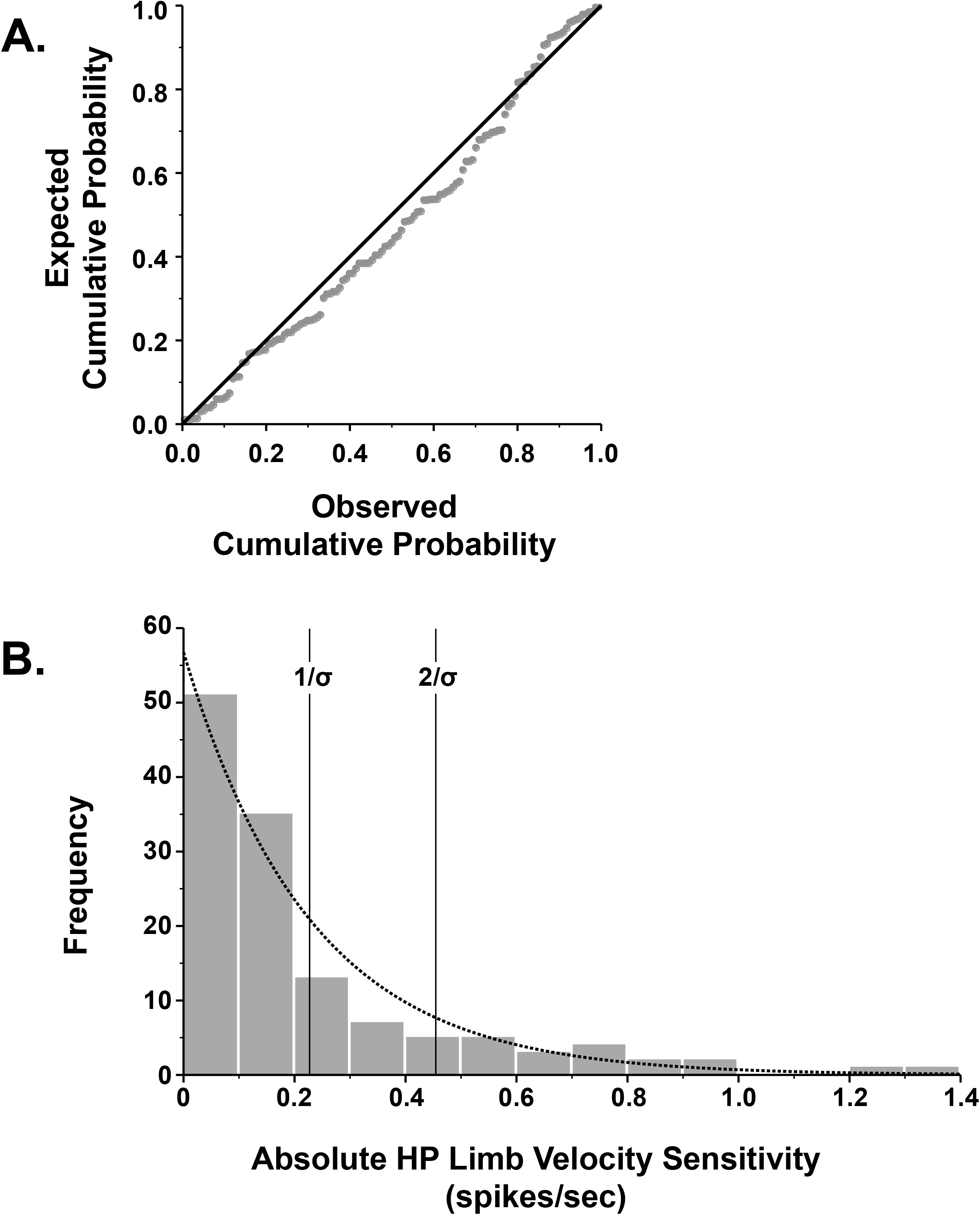
**A.** Exponential P-P plot of the absolute value of the HP limb velocity sensitivity variable. **B.** A histogram of the distribution of the absolute value of the limb sensitivity variable is positively skewed and is fit-best by an exponential distribution with a scale parameter (σ) of 4.401. The units were divided into distinct classes on the basis of the reciprocal of the scale parameter. Units were classified as having either low absolute HP limb velocity sensitivity, intermediate absolute HP limb velocity sensitivity, or high absolute HP limb velocity sensitivity.

The majority of neurons were classified into the low sensitivity to HP limb velocity category (69.8%, 90/129 neurons) and had sensitivities ranging from 0.002 to 0.227 spikes/sec (mean ± SD, 0.092 ± 0.064 spikes/sec). Twenty neurons (20/129 neurons; 15.5%) were classified as having intermediate HP limb velocity sensitivity. The mean sensitivity for this group was 0.327 ± 0.067 spikes/sec and values ranged from a low of 0.245 to a high of 0.438 spikes/sec. The remaining 19 neurons (19/129 neurons; 14.7%) were classified as having high absolute sensitivity. The mean sensitivity was 8.3 times higher than that for low sensitivity neurons (mean ± SD, 0.764 ± 0.238 spikes/sec; range, 0.475 - 1.371 spikes/sec).

### 3.5. Combined Vestibular and HP Limb Velocity Sensitivities

As with the combined vestibular and LP limb position sensitivities, neurons could be further divided into nine groups on the basis of vestibular and HP limb velocity sensitivities (Fig. 10A; Table 3). The most common unit responses (57/129 units; 44.2%) were neurons with low absolute vestibular - low absolute HP limb velocity sensitivities (Fig. 10B1). Intermediate absolute vestibular – low absolute HP limb velocity sensitivity neurons (20/129 units; 15.5%; Fig. 10B4) and low absolute vestibular – high absolute HP limb velocity sensitivity neurons (16/129, 12.4%; Fig. 10B3) were the next most common subcategories. High absolute vestibular – low absolute HP limb velocity sensitivity neurons were robustly sensitive to vestibular stimulation but were minimally responsive to limb movement made up 10.1% (13/129 neurons; Fig. 10B7) of the population. Low absolute vestibular – intermediate absolute HP limb velocity sensitivity neurons made up seven percent (9/129 neurons; Fig. 10B2). Intermediate absolute vestibular – intermediate absolute HP limb velocity sensitivity neurons made up six percent of the population (6/129 neurons; Fig. 10B5). A single neuron had intermediate absolute vestibular - high absolute HP limb sensitivity (0.8%; Fig. 10B6). Five neurons (3.9%) were highly sensitivity to vestibular stimulation and had intermediate sensitivity to absolute HP limb velocity (Fig. 10B8). Finally, two neurons were classified as having high absolute vestibular - high absolute HP limb velocity sensitivity (1.6%; Fig. 10B9).

**Figure 10.**
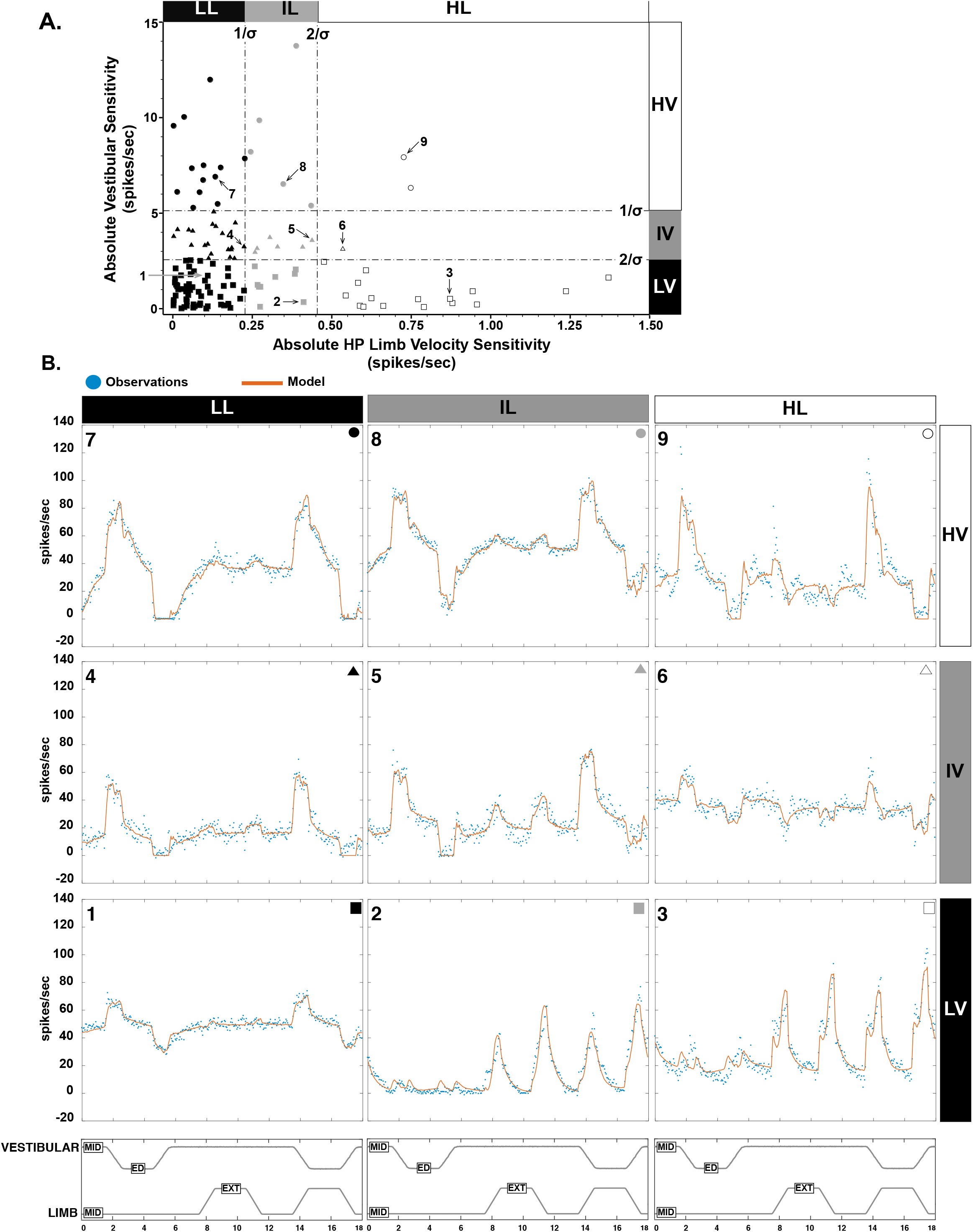
**A.** Neurons could be further divided into nine groups on the basis of vestibular and high pass limb velocity sensitivities. Units labeled 1 through 9 correspond to the example units shown in B. **B.** The averaged responses of 9 units represent the salient features of VN neuron responses to combined vestibular and hindlimb stimulation. The observed firing rate (blue dots) and the modeled firing rate (the fit of the model to the data; orange solid line, see section 3.4) are shown for each unit. Units 7-9 are classified as having high absolute vestibular sensitivity, units 4-6 have intermediate absolute vestibular sensitivity, and units 1-3 have low absolute vestibular sensitivity. Units 1,4, and 7 have low absolute high pass limb velocity sensitivity, units 2, 5, and 8 have intermediate absolute high pass limb velocity sensitivity, while units 3, 6, and 9 have high absolute high pass limb velocity sensitivity. Abbreviations: *ED, ear down; EXT, extension; MID, midline*

**Table 3.**

| **Table 3** |  |  | Absolute Vestibular Sensitivity (spikes/sec) |  |  |  | Absolute HP Limb Velocity Sensitivity (spikes/sec) |  |  |  |
| --- | --- | --- | --- | --- | --- | --- | --- | --- | --- | --- |
| **Class** | **%** | **Count** | **Mean** | **SD** | **Min** | **Max** | **Mean** | **SD** | **Min** | **Max** |
| LL | 44.20% | 57 | 0.969 | 0.742 | 0.006 | 2.538 | 0.079 | 0.061 | 0.002 | 0.224 |
| LI | 7.00% | 9 | 1.302 | 0.726 | 0.097 | 2.213 | 0.317 | 0.061 | 0.258 | 0.412 |
| LH | 12.40% | 16 | 0.784 | 0.726 | 0.088 | 2.455 | 0.782 | 0.253 | 0.475 | 1.371 |
| IL | 15.50% | 20 | 3.586 | 0.701 | 2.625 | 5.067 | 0.126 | 0.063 | 0.003 | 0.224 |
| II | 4.70% | 6 | 3.310 | 0.287 | 2.950 | 3.726 | 0.335 | 0.074 | 0.259 | 0.438 |
| IH | 0.80% | 1 | 3.122 | - | 3.122 | 3.122 | 0.535 | - | 0.535 | 0.535 |
| HL | 10.10% | 13 | 7.561 | 1.932 | 5.284 | 11.987 | 0.094 | 0.062 | 0.003 | 0.227 |
| HI | 3.90% | 5 | 8.747 | 3.271 | 5.392 | 13.755 | 0.338 | 0.079 | 0.245 | 0.435 |
| HH | 1.60% | 2 | 7.128 | 1.135 | 6.325 | 7.930 | 0.737 | 0.016 | 0.726 | 0.748 |

**Abbreviations:** LL, Low vestibular-low limb; LI, Low vestibular-intermediate limb; LH, Low vestibular-high limb; IL, Intermediate vestibular-low limb; II, Intermediate vestibular-intermediate limb; IH, Intermediate vestibular-high limb; HL, High vestibular-low limb; HI, High vestibular-intermediate limb; HH, High vestibular-high limb

### 3.6. Baseline Firing Rate

The estimated baseline firing rates of vestibular nucleus neurons ranged from −34 to 116 spikes/sec (mean ± SD, 24 ± 20 spikes/sec). The distribution of baseline firing rates is illustrated in figure 11. Approximately seven percent (9/129) of neurons had a zero or negative estimated baseline firing rate. Neurons with a zero or negative baseline firing rate were silent and responded only during vestibular or limb stimulation, suggesting that they were under tonic inhibition (e.g. Figs. 8B3 and 10B2). The remaining 120 units had a positive estimated baseline firing rate (e.g. Fig. 10B1). Like vestibular afferents, a positive baseline firing rate permits the neurons to display directional tuning. The baseline firing rate did not appear to differ as a function of either the magnitude of the absolute vestibular sensitivity (Fig. 12A), the magnitude of the absolute LP limb position sensitivity (Fig. 12B), or the magnitude of the absolute HP limb velocity sensitivity (Fig. 12C).

**Figure 11.**
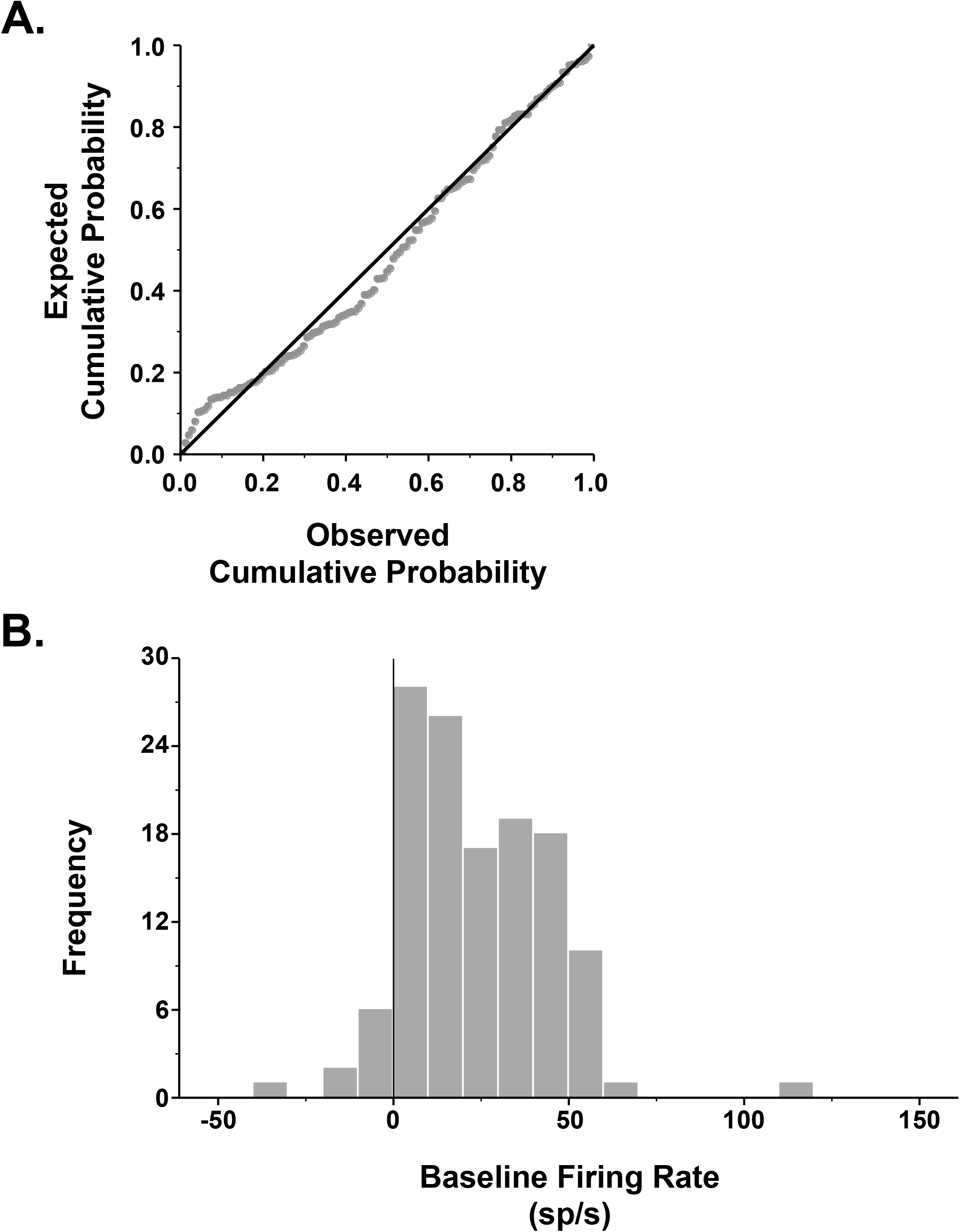
Modeled baseline firing rate of vestibular nucleus neurons. **A.** Normal P-P plot of the baseline firing rate. **B.** A histogram of the distribution of the baseline firing rate is fit-best by a normal distribution with mean of 24 and standard deviation of 20 (spikes/sec).

**Figure 12.**
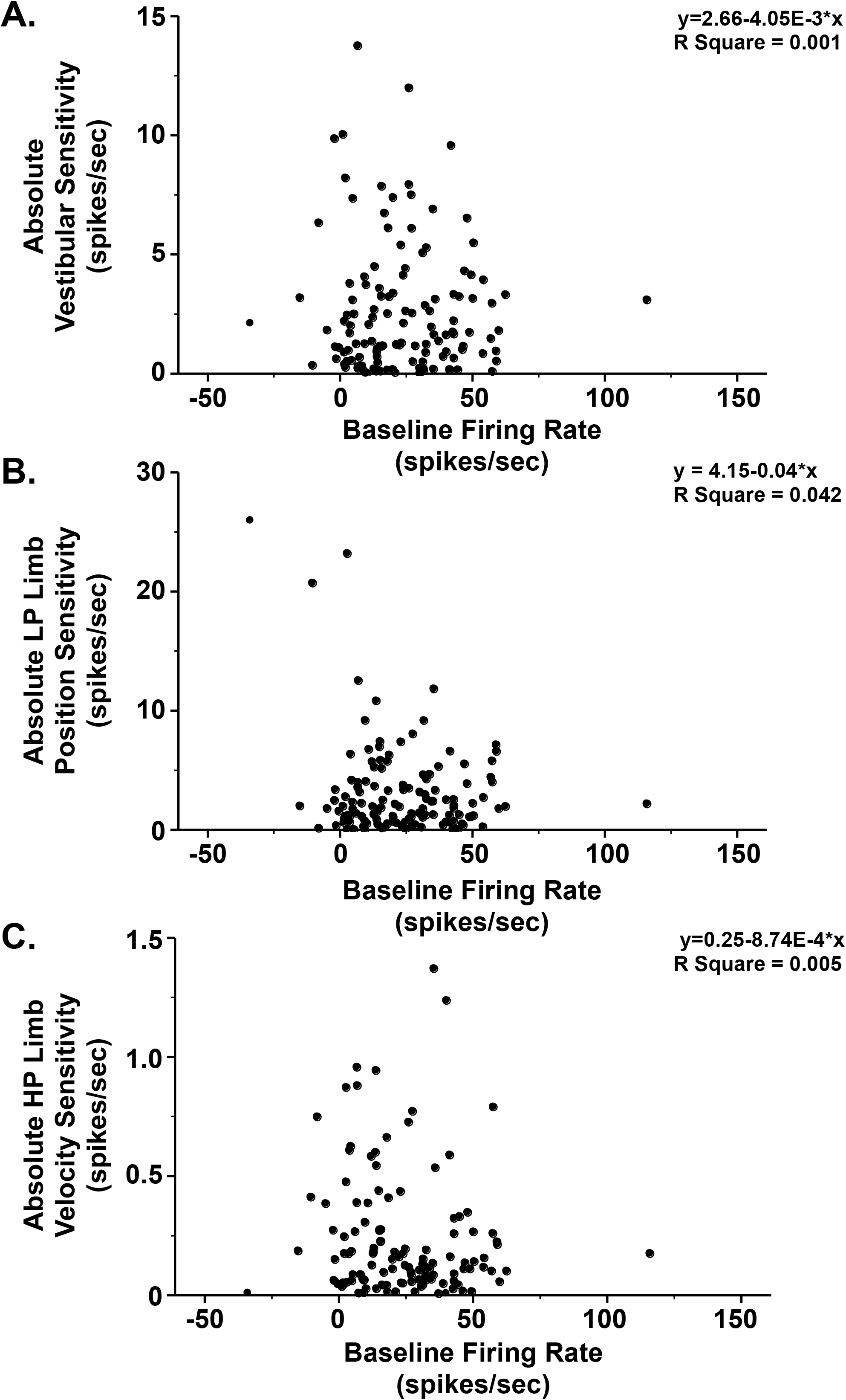
The baseline firing rate was not correlated with the (A) absolute vestibular sensitivity, (B) absolute LP limb position sensitivity, (C) or absolute HP limb velocity sensitivity. The regression equation and R^2^ value for each subplot are indicated in the upper right of the subplot.

### 3.7. Location of VN Units

The physiological data were obtained from 129 units recorded from the brainstem of four conscious felines. The locations of recorded neurons were plotted with respect to an electrolytic lesion made in the same tract or in an adjacent track. It was possible to accurately reconstruct the location of 88 neurons. These 88 neurons were histologically confirmed to be located within the lateral, medial, and inferior vestibular nuclei (Fig. 13). Units were scattered over a rostrocaudal extent of 2.1 to 7.5 mm (mean 5.5 ± 1.8 mm) rostral to the obex and were on average, 2.6 ± 1.0 mm lateral to the midline (range: 0.5 to 5.3 mm). Lesions were not well formed in one animal (accounting for 41 neurons in the dataset), we thus relied on scarring from recording tracks to plot out the locations of recorded units. In this animal, units were located over of a similar spread of territory (distance rostral to the obex: 3.3 to 8.1 mm (5.8 ± 1.1 mm); distance lateral to the midline: 1 to 4 mm (3.0 ± 0.8 mm)) as the other three animals. Because we were not able to definitively confirm locations within the specific vestibular subnuclei relative to lesion sites in this animal, these data are not plotted on Fig. 13.

**Figure 13.**
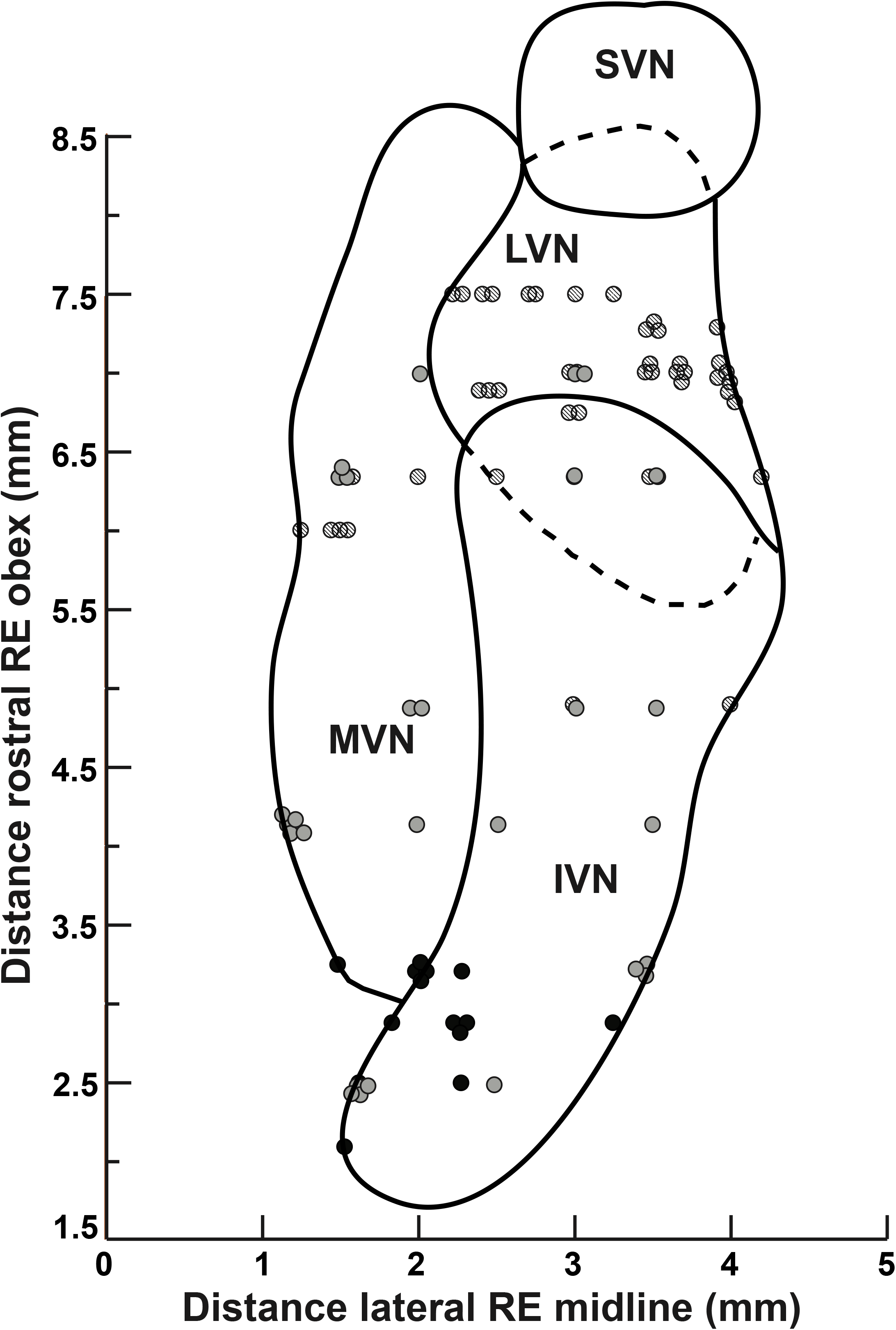
A horizontal section through the vestibular nucleus complex showing the location of neurons whose activity was recorded in this study. The y-axis designates the distance in millimeters rostral to the obex and the distance in millimeters lateral to the midline is shown on the x-axis. The activity of neurons was recorded over a rostral-caudal distance of 6 mm, from 2.1 mm to 8.1 mm rostral to the obex. Symbol shading indicates neurons recorded from different cats. Abbreviations: IVN, *inferior vestibular nucleus;* LVN, *lateral vestibular nucleus;* MVN, *medial vestibular nucleus;* RE, *relative to; SVN superior vestibular nucleus*.

## 4. Discussion

Our findings demonstrate that vestibular nucleus neurons integrate information from hindlimb proprioceptive inputs with information from peripheral vestibular receptors. We found that VN neuronal activity during combined vestibular and hindlimb proprioceptive stimulation in the conscious cat is well-fit by a simple additive model for signals with similar temporal dynamics. The average R^2^ goodness-of-fit value for fitting the additive model to neural firing was 0.74, with very few model fits being in the lower range (Fig. 5). The ability to integrate multimodal sensory signals is a key feature of neurons in the central nervous system that participate in balance control and is particularly notable among those in the vestibular nuclei [1–4, 8, 10, 33]. Convergence of proprioceptive signals with vestibular signals commonly occurs among vestibular nucleus neurons. For instance, in cats and squirrel monkeys a substantial majority (~80%) of vestibular nucleus neurons were found to have their activity modulated with activation of neck proprioceptive afferents and those neck proprioceptive signals integrated with vestibular inputs in an additive fashion [25, 26, 34–37]. Our group has previously demonstrated that over two-thirds of vestibular nucleus neurons in conscious cats have their activity modulated by hindlimb movement [17]. The present work extends those previous findings by demonstrating that limb proprioceptive signals integrate with vestibular signals in an additive fashion. Since proprioceptive inputs from the neck and hindlimb each converge and integrate with vestibular inputs in the vestibular nuclei in an additive manner, it is likely that proprioceptive inputs from throughout the body (including, for example, inputs from proprioceptors signaling movement using the thoracic and lumbar axial musculature and from the forelimbs, in quadrupeds at least) are similarly processed by the vestibular nuclei. This suggests that vestibular nucleus neurons may process vestibular afferent information in the context of position and movement of the whole-body in space, rather than solely in a “head-referenced” frame [38, 39]. This is likely especially true of vestibular nucleus neurons with spinal projections that influence motor outflow in response to perturbations.

While our final model included eight vestibular and limb related signal components and one baseline component, we chose to focus our analysis on the dynamics from three main components: vestibular sensitivity (rectified HP roll position sensitivity + 0.291* HP roll velocity sensitivity (ipsilateral ear-up positive)), LP limb position sensitivity (rectified LP limb position sensitivity - 5.68 sp/s (a constant to adjust its set point)) and HP limb velocity sensitivity (HP limb flexion velocity sensitivity – 0.402*HP limb extension velocity sensitivity) because these terms included features that were thought to be most representative of the underlying stimuli.

The vestibular sensitivity term selected for in-depth analysis is thought to represent a processed signal containing inputs originating from both semicircular canal and otolith afferents. Roll plane stimulation was chosen because it has previously been demonstrated that the vast majority of neurons receiving converging vestibular and hindlimb inputs have response vector orientations within 45° of the roll plane [16, 17]. Semicircular canal and dynamic otolith afferents likely drive the high pass roll velocity signal. The semicircular canals have high-pass filtering properties, and due to the fluid mechanics of the canals, the response of canal afferents is in phase with the velocity of rotation [40]. While semicircular canals have dynamic components, otolith units have both dynamic and static components [41]. Phasic-tonic otolith afferents have a dynamic component that scales with increasing ramp velocity and a static positional component during the hold period [41]. The rectified high pass roll position signal is likely a highly processed otolith signal. Evidence from non-human primates shows that there is substantial spatiotemporal processing of otolith afferent signals by vestibular nucleus neurons [42]. In the case of the limb terms, selecting representative terms was moderately more challenging because, unlike vestibular inputs, little is presently known about which specific proprioceptive receptors (muscle spindles, cutaneous receptors, Golgi tendon organs, etc.) are responsible for signaling limb-state changes to the vestibular nuclei. However, responses of vestibular nucleus neurons in the conscious cat have a strong phasic component during limb movement and, in some cases, a lasting positional component [17], which are well approximated by the HP limb velocity term and LP limb position term, respectively.

Sensitivities of vestibular nucleus neurons to individual components of the additive model took the form of exponential functions (see Figs. 6, 7, and 9), excepting estimated baseline firing rate (see Fig. 11), with the majority of neurons exhibiting low sensitivity to a given component and relatively few being highly sensitive. Interestingly, neurons with high sensitivity to one domain were unlikely to also have high sensitivity in a cross-modal domain. For example, as shown in Fig. 8, most neurons that were in the high category of LP limb position sensitivity were in the low category of vestibular sensitivity and vice versa. These findings suggest that while a large majority of the population of vestibular nucleus neurons appear fairly broadly tuned to cross-modality stimuli, some subpopulations of neurons appear to be tuned to extract particular salient features of the stimuli in one modality. When viewed as a population of neurons that is attempted to discern input characteristics of a given stimulus, having neurons that excel at encoding particular features of the stimulus could allow for subcortical sensory processing of salient features leading to balance percepts. Note, however, that while seemingly useful, having distinct response properties to salient features of a stimulus is not a necessary feature for subcortical sensory perceptual processing. Jorntell and others [33] recently demonstrated that salient features of skin-object interactions were encoded by neurons in the cuneate nucleus despite having similar underlying response properties and receptive fields.

It is important to remember that the data generated in this study were all from passively applied limb movements, and thus the responses elicited in the vestibular nuclei were in response to exafferent signals. Because of this fact, one reasonable hypothesis to consider is that the information encoded by VN neurons in response to passive limb movement serves to prime vestibulospinal reflexes during destabilizing postural occurrences, in particular those that result in co-activation of vestibular and limb afferents. An example of such a situation where this may be beneficial is a person walking on a sidewalk and unexpectedly encountering black ice. In such a situation, when the lower limb slips a limb exafferent signal would be generated concurrent with an exafferent signal from vestibular afferents due to the co-occurring resultant head movement in space. Such a situation would call for rapid postural adjustments in an attempt to maintain upright stance, and as such, a primed vestibulospinal response would be warranted. Active (self-generated) hindlimb muscle contraction can also modulate the activity of vestibular nucleus neurons [17], however it remains unclear if the resultant modulation under those conditions is from reafference or efference copy. As the present study did not examine for integration of limb proprioceptive signals with vestibular signals under conditions of active movement, it is presently unknown how these signals would influence vestibular nucleus neuronal activity during combined volitional limb and head movement, such as happens during locomotion [28]. It is possible that integration of these signals would vary substantially during conditions of active movement, as has been seen in other studies examining the influence of signals from semicircular canals and otoliths on vestibular nucleus neuronal activity. During conditions of combined translation and rotation in non-human primates, the combined integrated signal is less during self-generated active movement than when these stimuli are passively applied [43]. Future studies will be needed to determine how strongly signals are integrated in the vestibular nucleus during combined active limb and head movements.

One limitation of the current study is that felines are quadrupeds and it is likely that inputs from all four limbs are relayed to vestibular nucleus neurons, whereas our setup focused on inputs from only one limb. Vestibulospinal neuronal activity has previously been shown to modulate with the step-cycle in conscious cats in which all four limbs are in motion [44, 45]. This finding raises the intriguing possibility that inputs from certain parts of the limbs, such as afferents innervating flexor or extensor muscle spindles, joint receptors, or cutaneous receptors may predominate at times depending upon movement about specific joints or activation of specific muscle groups with the step cycle. However, those experiments were carried out in head-free animals and thus the relative contribution of vestibular inputs was not able to be controlled for. Additional experiments are needed to explore how inputs from all four limbs integrate in the vestibular nuclei, particularly in head-fixed animals, under active and passive conditions. Another limitation of our experimental setup is that we did not define the projections of the vestibular nucleus neurons that we were recording from. It is likely that a large proportion of these neurons are vestibulospinal neurons that descend via the spinal cord to indirectly and directly influence motoneuron activity. Evidence from human studies shows that vestibulospinal reflexes are altered to account for lower limb [46, 47] and head position [1, 3, 9, 48–50]; the neuronal responses we observed suggest vestibular nucleus neurons play a key role in mediating these responses. It is also possible that output information from vestibular nucleus neurons that integrate vestibular and limb proprioceptive information could influence other systems such as mediation of vestibuloautonomic reflexes [51], motor planning and execution through cerebellar projections, or project to cortical regions via vestibulothalamic pathways [52].

In conclusion, neurons in the feline VN integrate inputs from the limb and labyrinth in an additive manner. We speculate that integration of these vestibular and limb signals by VN neurons serve to adjust vestibulospinal reflexes to account for limb position in space when a balance perturbation occurs. Further research will be needed to determine if this population of neurons has descending efferent projections down the spinal cord. This knowledge of how single neurons integrate sensory information across modalities is crucial to understanding how the vestibular system uses multimodal information to maintain balance in a dynamic environment.

## Acknowledgements

This work was supported by NIH funding from F32-DC015157 (Derek M. Miller) and K08-DC013571 (Andrew A. McCall).

